# CD38 regulates ovarian function and fecundity via NAD^+^ metabolism

**DOI:** 10.1101/2023.05.08.539779

**Authors:** Rosalba Perrone, Prasanna Vadhana Ashok Kumaar, Lauren Haky, Cosmo Hahn, Rebeccah Riley, Julia Balough, Giuliana Zaza, Bikem Soygur, Kaitlyn Hung, Leandro Prado, Herbert G. Kasler, Ritesh Tiwari, Hiroyuki Matsui, Genesis Vega Hormazabal, Francesca Elizabeth Duncan, Eric Verdin

## Abstract

Mammalian female reproductive lifespan is typically significantly shorter than life expectancy and is associated with a decrease in ovarian NAD+ levels. However, the mechanisms underlying this loss of ovarian NAD+ are unclear. Here, we show that CD38, a NAD+ consuming enzyme, is expressed in the ovarian extrafollicular space, primarily in immune cells, and its levels increase with reproductive age. Reproductively young mice lacking CD38 exhibit larger primordial follicle pools, elevated ovarian NAD+ levels, and increased fecundity relative to wild type controls. This larger ovarian reserve results from a prolonged window of follicle formation during early development. However, the beneficial effect of CD38 loss on reproductive function is not maintained at advanced age. Our results demonstrate a novel role of CD38 in regulating ovarian NAD+ metabolism and establishing the ovarian reserve, a critical process that dictates a female’s reproductive lifespan.

## Introduction

The ovary is a dynamic organ consisting of heterogeneous cell types that must function coordinately to enable female reproductive function. In mammals, the ovarian reserve or number of primordial follicles that dictates the reproductive lifespan is established early in development and is considered finite and non-renewable. The ovarian reserve undergoes a natural winnowing over time combined with diminished egg quality, which together contribute to decreased fertility in females by their late 30s and eventual complete cessation of reproductive function at menopause ^1^.

Cells within the ovary require energy to maintain homeostasis to support folliculogenesis ^2^. Nicotinamide Adenine Dinucleotide (NAD+) is a cofactor, co-substrate, and redox partner in important metabolic pathways involved in energy generation, cell signaling, and tissue repair ^3^. NAD+ regulates energy metabolism in several ovarian cells, including oocytes, granulosa cells, and theca cells, which orchestrate folliculogenesis, oocyte maturation, and ovulation ^4, 5^. During ovulation, there is a high demand for NAD+ to support follicular rupture, cumulus-oocyte complex release, and corpus luteum formation. NAD+ also contributes to DNA damage repair which is critical for the prevention of chromosomal abnormalities and maintenance of genetic integrity ^6^. Considering the central role of NAD+ in mediating ovarian metabolism and henceforth function, it is not surprising that there is a strong association between a decline in ovarian NAD+ levels and loss of ovarian function.

NAD+ homeostasis is maintained by a ‘supply-demand’ chain, in which NAD+ is replenished by biosynthetic pathways to counter its utilization by NAD+-dependent metabolic pathways ^2, 3^. However, over time, this balance tips towards the latter due to increased demand for NAD+ under survival circumstances, such as the constant need for DNA repair caused by cumulative oxidative stress ^7, 8^. The decline in NAD+ with chronological aging has been reported in several organs, such as the liver and adipose, and is detrimental to physiological processes leading to metabolic dysfunction and age-associated diseases ^9^. In the ovary, NAD+ levels also decline with age, and this reduction is associated with a decrease in gamete quantity and quality ^10–12^. Supplementation with NAD+ precursors, such as nicotinamide mononucleotide (NMN) or nicotinamide riboside (NR), increase NAD+ levels and improve gamete quality, ovarian function, and fertility in aged mice ^10–13^. These findings suggest a key role for NAD+ metabolism in ovarian function and reproductive aging. However, the molecular mechanisms regulating ovarian NAD+ loss have not been elucidated.

The age-related decrease in NAD+ levels is not solely attributed to an increased demand for NAD+ but also to an increase in NAD+ degradation ^14^. One of the potential factors contributing to the NAD+ deficit during aging is the glycoprotein NADase, CD38, which catalyzes the breakdown of NAD+ into nicotinamide (NAM) and ADP-ribose (ADPR) ^15, 16^. CD38 mediates key cellular functions, from regulating immune cell function as a surface receptor on specific innate and acquired immune cells to modulating metabolic processes through its hydrolase and cyclase activities ^17, 18^. However, an age-associated increase in CD38 expression and activity in multiple tissues, such as the liver, adipose, and skeletal muscle, negatively correlates with tissue NAD+ levels, corroborating its role in age-associated cellular dysfunction ^19, 20^. Although there is existing research on CD38’s role in aging and age-related diseases ^21–23^, its role in female reproductive function and aging has not been systematically investigated.

In the current study, we demonstrate that CD38 is expressed in the mouse ovary primarily in the extrafollicular compartment with an enrichment in immune cells. Ovarian CD38 increases with age, and interestingly, our findings using a genetic loss-of-function model demonstrate that CD38 is a negative regulator of reproductive function in reproductively adult females. Female mice lacking CD38 exhibit a larger number of ovarian follicles and increased fecundity up to 5 months of age, and these phenotypes are associated with higher ovarian NAD+ levels. The increased number of follicles appears to be due to the role of CD38 in modulating ovarian reserve formation early in development. Overall, our findings highlight the critical role of CD38 in regulating ovarian function and fertility through NAD+ metabolism. Targeting CD38 or its downstream effects on NAD+ metabolism may offer new approaches to enhance reproductive longevity.

## Results

### 1. CD38 is expressed in extrafollicular cells in the ovary and its expression increases with age

The ovary is a heterogeneous tissue that is constantly subject to remodeling due to folliculogenesis, ovulation, and corpus luteum formation and regression ^24^. The ovary includes multiple cell types, including granulosa, mesenchymal, endothelial, epithelial, immune cells, and oocytes, and the specific composition is highly dependent on animal age and cycle stage ^25^.

To determine where CD38 is expressed in the ovary, we visualized and quantified *Cd38* transcripts in specific ovarian subcompartments in reproductively adult mice (Figure 1A). Cells within extrafollicular subcompartments expressed high levels of *Cd38* transcript, including the ovarian surface epithelium (OSE), the stroma, the vasculature, and the corpora lutea (CL) (Figure 1B). Interestingly, follicles at all stages of development exhibited minimal to no *Cd38* expression both in the oocyte and somatic granulosa cells (Figure 1C). Computational image analysis in which *Cd38* mRNA transcript number in each subcompartment was normalized to subcompartment area confirmed that ovarian *Cd38* expression was primarily extrafollicular with CL and vascular compartments having the highest levels of *Cd38* transcription when compared to OSE and ovarian follicles (Figure 1D).

**Figure 1:**
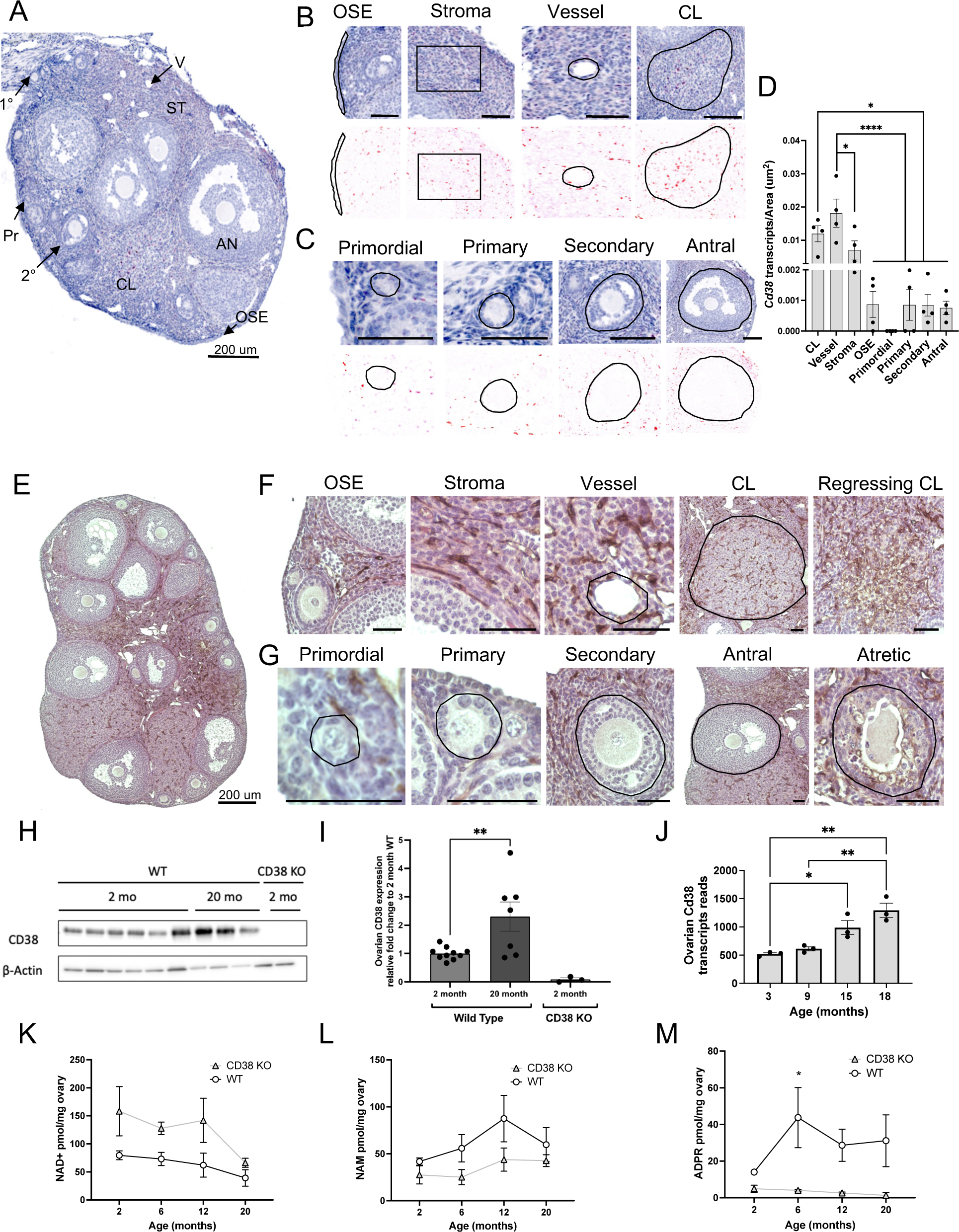
CD38 is expressed in the extrafollicular regions of the mouse ovary, its expression increases with age, and it regulates ovarian NAD+ metabolism. (A) Localization of CD38 mRNA by in situ hybridization (RNAscope) in 2-month-old ovarian section from WT mouse, 200 µ scale. (B) Higher magnification images from Figure 1A showing CD38 mRNA localization by RNA Scope in OSE, stroma, vessel and CL. 50 **Mµµµ**scale unless otherwise indicated. (C) Higher magnification images from Figure 1A showing CD38 mRNA localization by RNA Scope in ovarian follicles at different developmental stages. 50 µm scale unless otherwise indicated. (D) Quantification of CD38 mRNA localization by RNA Scope in ovarian structures and regions (n=4). (One-Way ANOVA statistical analysis, * = p < 0.05, **** = p < 0.00005). (E) Localization of CD38 protein by immunohistochemistry (IHC) in 2-month-old ovarian section from WT mouse, 200 µm scale (F) Higher magnification images from figure 1E showing CD38 protein localization in OSE, stroma, vessel and CL and regressing CL, 50 µm scale unless otherwise indicated (G) Higher magnification images of CD38 mRNA localization by IHC in ovarian follicles at different developmental stages, 50 µm scale unless otherwise indicated. (H) Representative Western blot of CD38 expression in whole ovarian tissue lysate from 2 months (n=6) and 20 months (n=3) old WT mice. Whole ovarian tissue lysate from 2 months old CD38 KO animals (n=2) was used as control. β-actin was used as housekeeping loading control. (I) Bar chart from 3 independent Western Blot experiments showing CD38 expression in whole ovarian tissue lysate from 2 months and 20 months old WT mice. Whole ovarian tissue lysate from 2 months old CD38 KO animals was used as control. (Unpaired T Test statistical analysis, **= p<0.005). (J) Cd38 transcript reads in 3, 9, 15, and 18 month old mouse ovaries (n=3). (One-Way ANOVA statistical analysis, *= p<0.05, **= p<0.005). WT and CD38 KO ovarian (K) NAD+, (L) NAM, and (M) ADPR levels at different ages quantified using LCMS (n=2-5 per group, 2-way ANOVA analysis, * = p <0.05). (OSE= ovarian surface epithelium, CL=corpus luteum)

CD38 protein expression largely paralleled the transcript (Figure 1E), localizing mainly to the ovarian stroma, the vasculature, the CL, the regressing CL, and the atretic follicles (Figure 1F). There was minimal to no expression in the OSE (Figure 1F). Follicles at all stages of development lacked CD38 protein expression in both the germ cell and granulosa cell compartments, but there appeared to be expression in the vascular region of the theca layer. Overall, CD38 protein localization was consistent with what was observed with the transcript (Figure 1G).

CD38 expression increases with age in several mouse tissues, including liver, adipose, spleen and skeletal muscle ^26^. To investigate how CD38 levels change with age in ovaries, we extracted whole tissue lysate from reproductively young (2 months) and old (20 months) mouse ovaries and evaluated total CD38 protein content via Western blot. We observed a more than 2-fold increase in CD38 protein levels with age (Figure 1H) (Figure 1I). These findings are consistent with a previously published transcriptomic dataset in which an increase in *Cd38* transcript levels is observed in mouse ovaries at 18 months of age ^27^ (Figure 1J).

### 2. CD38 KO animals have increased ovarian NAD+ levels and decreased NAM and ADPR levels

An age-dependent increase in CD38 expression causes NAD+ level depletion in multiple tissues, and a total body knock-out mouse model of CD38 (CD38 KO) is protected from NAD+ loss ^26^. Although previous research has documented a decline in ovarian NAD+ levels ^10, 11^, the role of CD38 in this process is unknown. Therefore, we investigated the impact of CD38 on ovarian NAD+ regulation. To this end, we collected ovaries from WT and CD38 KO females at different reproductive ages (2, 6, 12 months) and post-fertile age (20 months) and measured NAD+ and related metabolites (NAM and ADPR) using mass spectrometry. Ovaries from CD38-deficient mice exhibited higher NAD+ levels than ovaries from age-matched WT mice across reproductive ages (2-12 months) (Figure 1K). However, at 20 months of age, NAD+ levels in ovaries from CD38 KO mice declined to levels comparable to those of WT controls, indicating that CD38 deficiency did not offer protection from NAD+ loss during physiologic ovarian aging (Figure 1K). Notably, ovaries from CD38 KO mice had decreased levels of NAM (Figure 1L) and ADPR (Figure 1M) relative to age-matched WT controls. These are both primary products of CD38-dependent NAD+ hydrolysis, thus highlighting the specific effect of CD38 on ovarian NAD+ metabolism. Our findings demonstrate that CD38 plays a crucial role in regulating and maintaining NAD+ homeostasis in ovaries during reproductive aging.

### 3. CD38 is enriched in ovarian immune cells and its expression in leukocytes increases with reproductive age

CD38 transcript and protein were highly expressed in the ovarian stroma which is heterogeneous in cell type composition, containing populations including fibroblasts, immune cells, endothelial cells, and smooth muscle cells ^28^. To gain a better understanding of cell type-specific *Cd38* expression in the ovary, we interrogated the Mouse Cell Atlas (MCA) which contains transcriptomic data of mouse tissues at single-cell resolution ^29^. *Cd38* was highly expressed in ovarian endothelial cells and macrophages ^29^, which is consistent with localization in other tissues^22^. To further examine CD38 expression in ovarian endothelial cells and macrophages, we performed co-labeling in ovarian histological sections with antibodies against CD38 and either CD31 (endothelial cell marker) or F4/80 (macrophage cell marker) Fig S1A-B). CD31 expression was enriched in some CLs and in the ovarian stroma as expected given the known localization of vasculature in the ovary (Fig S1 A). Only a portion of CD31 expressing cells co-labeled with CD38 within the ovarian stroma and vessels (Fig 2A’). Interestingly, there were distinct populations of both CD38 positive (Fig S1A’’) and negative (Fig S1A’’’) endothelial cells within the corpora lutea. Similar to CD31, F4/80 positive cells were mostly localized in the ovarian stroma and the CLs (Fig 2B). The majority of F4/80 expressing cells appeared to be CD38 positive within the ovarian stroma (Fig S2B’) and CD38 negative in the corpora lutea (Fig S2B’’), suggesting heterogeneity of CD38 expression within the ovarian macrophage population. Our results confirmed that CD38 is expressed in a subset of ovarian endothelial cells and macrophages but also demonstrates localization to other cell types within the ovarian microenvironment.

**Figure 2:**
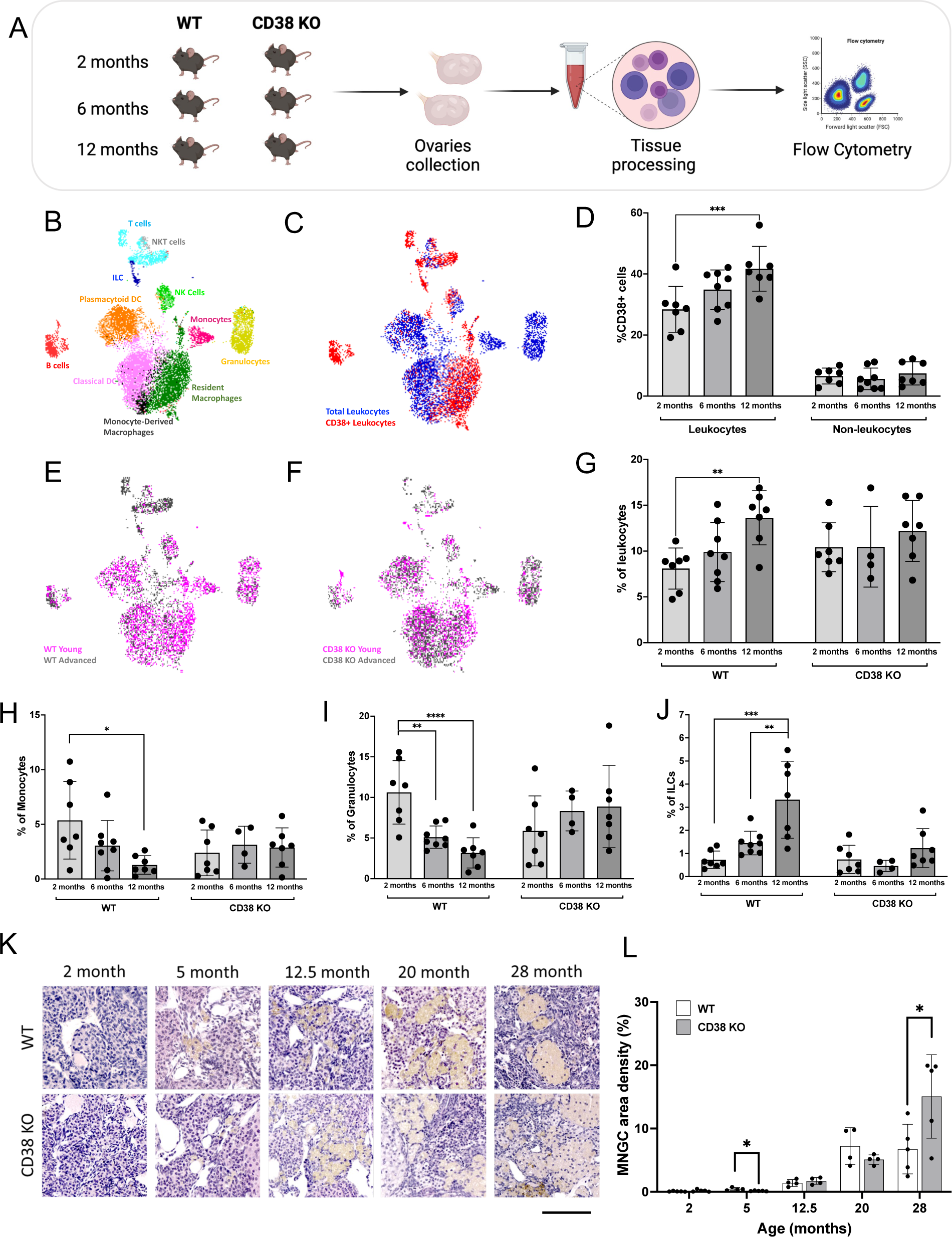
CD38 expressing ovarian immune cells increase with age and CD38 KO animals are partially protected from age-associated ovarian immune changes. (A) Schematic of Flow Cytometry-based ovarian immunophenotypic analysis of 2, 6 and 12 months old females. (B) UMAP plot featuring the different immune cell subclusters belonging to the murine ovarian immune landscape (C) UMAP plot featuring CD38 expressing immune cells within the WT ovarian immune landscape (D) Percentage of CD38 positive (CD38+) cells within the leukocytes (CD45+) and non-leukocytes (CD45-) population throughout murine reproductive aging (n=7-8 per group, mean ± SEM, Two-way ANOVA statistical analysis *** = p < 0.0005). UMAP plot featuring immune cells distribution within the young and old (E) WT and (F) CD38 KO ovarian immune landscape. (G) Percentage of total leukocytes (CD45+) population throughout murine reproductive aging in WT and CD38 KO ovaries (n=7-8 per group, mean ± SEM, Two-way ANOVA statistical analysis ** = p < 0.005). Percentage of total (H) Monocytes (CD11b+, F4/80-, Ly6C+, Ly6G-) (I) Granulocytes (CD11b+, F4/80-, Ly6C+, Ly6G+) and (J) ILCs (CD3-, CD19-, CD138-, NK1.1-, CD11b-, CD127+) throughout murine reproductive aging in WT and CD38 KO ovaries (n=4-8 per group, mean ± SEM, Two-way ANOVA statistical analysis * = p < 0.05, ** = p < 0.005, *** = p < 0.0005, **** = p < 0.00005). (K) Representative images of H&E stained ovarian sections from 2, 5, 12.5, 20, and 28 month old WT and CD38KO mice visualized by bright-field microscopy, 100 µm scale. (L) Quantification of MNGCs density in ovarian sections from 2, 5, 12.5, 20, and 28 month old WT and CD38KO females (n=5 per group) (Unpaired t-test statistical analysis, * p< 0.05).

In addition to macrophages, there are many different populations of immune cells in the ovary that undergo dynamic changes during development, the reproductive cycle, and aging ^27, 30^. In other tissues, CD38 is expressed in several immune populations, including macrophages, NK, T and B cells ^31^. CD38 expression on immune cells increases as a result of key immunological responses such as inflammation and infection ^31^, and during aging an accumulation of CD38 positive immune cells, in particular macrophages, has been reported in several tissues ^22, 23^ which drives NAD+ depletion and leads to age-associated phenotypes ^20, 26^.

To further characterize CD38 expression within the ovarian microenvironment during reproductive aging, we performed a flow cytometry-based immune profiling analysis of the mouse ovary across the reproductive lifespan at reproductively young, mid, and advanced ages. We analyzed ovaries from both WT and CD38 KO mice to capture potential cell composition changes in the absence of CD38 (Figure 2A). We identified eleven cell types representing the main populations involved in innate and adaptive immunity (Figure 2B). CD38 was expressed in different immune cell types, primarily macrophages, B cells, NK cells, and T cells (Figure 2C), in line with previously reported evidence in other tissue types ^31^. In ovaries from WT mice, approximately 30-40% of the cells expressing CD38 were within the ovarian leukocyte population (CD45+), whereas only approximately 10% of the non-leukocyte (CD45-) cells in the ovary expressed CD38 (Figure 2D). Interestingly, CD38+ leukocytes increased with age, whereas CD38+ cells in the non-leukocyte population did not change (Figure 2D). These data are consistent with our previous data in the liver where CD38 levels increased with age only in Kupffer cells but not in non-immune cells including hepatocytes and endothelial cells ^22^. Our results demonstrate that ovarian CD38 is mostly expressed in the immune compartment, and CD38 positive leukocytes specifically increase with reproductive age.

### 4. CD38 loss-of-function results in significant changes in the ovarian immune milieu during reproductive aging

Consistent with other reports ^27, 32^, our data demonstrated that the ovarian immune compartment profoundly shifts with advanced reproductive age (Figure 2E). Ovaries from reproductively young WT mice are enriched in innate immune cells, including macrophages and dendritic cells (DC) which together represent around 60% of the entire ovarian immune landscape (Figure S2A). Interestingly, ovaries from reproductively old mice exhibit a dramatic decrease in innate immunity coupled with an increase in adaptive immunity, in particular T cells (Figure S2A). The age-dependent increase in T cells appeared to be mostly driven by CD4,CD8 double negative (DN) T cells (Figure S2B).

Significant age-related changes in the immune milieu were also observed in ovaries from CD38 KO mice (Figure 2F) with overall similar trends in reduced innate immunity and increased adaptive immunity (Figure S2C). The main adaptive population that increased with age in ovaries from CD38 KO mice was the DN T cells similar to WT controls (Figure S2D). Although overall age-dependent trends in immune cell populations were similar across genotypes, CD38 KO mice showed several unique features compared to WT counterparts. Ovaries from wild type mice showed an age-dependent increase in total leukocytes while the number of immune cells in CD38 KO ovaries remained constant throughout reproductive aging (Figure 2G). Additionally, ovaries from WT mice showed an age-dependent decrease in monocytes (Figure 2H) and granulocytes (Figure 2I), but this pattern was absent in CD38 KO ovaries (Figure 2H-I), suggesting a role of CD38 in regulating ovarian innate immune composition during aging. Innate lymphoid cells (ILCs), another cell type of the innate immunity, significantly increased with age in ovaries from WT mice, but only modest changes in this population were observed with age in ovaries from CD38 KO mice (Figure 2J). As for other innate immune cells, an overall age-dependent decrease in total macrophages was observed in both WT and CD38 KO ovaries while no changes in DC occurred in both strains (Figure S2A-B). Interestingly, we noticed a change in the macrophage cell composition with a relative increase with age in the monocyte-derived population (CD11c+) versus the tissue-resident (CD11c-) population in both WT and CD38 KO ovaries (Figures S2E-F). The same age-related trend in ovarian macrophage composition was previously reported ^27^.

In addition to canonical macrophages, a highly penetrant and unique population of macrophage-derived multinucleated giant cells (MNGCs) accumulates with age in the ovary and are associated with chronic inflammation ^33, 34^. Although this cell population has been demonstrated to express the macrophage marker F4/80 ^34^, their analysis and quantification with single-cell based techniques, including Flow Cytometry, is challenging due to their large size and limited survival during cell-dissociation protocols. Histological analysis of MNGCs represents the optimal way to visualize and quantify MNGCs in tissues ^33, 35^. Given the importance of MNGCs in the context of ovarian aging, we analyzed their presence and relative abundance in ovaries from both CD38 KO and WT mice collected across ages spanning 2 to 28 months (Figure 2K and Figure S3A-B). As expected, MNGCs were virtually absent in ovaries from reproductively young mice and significantly increased with age (Figure 2L). No significant differences in the amount of MNGCs was observed between genotypes during reproductive ages (2 to 12 months). However, a significantly larger ovarian area was occupied by MNGCs in ovaries from CD38 KO mice at 28 months relative to controls (Figure 2L). Overall our data suggest an immuno-modulatory role of CD38 in shaping the ovarian immune landscape during reproductive aging.

### 5. CD38 loss-of-function results in a larger ovarian reserve and enhanced fecundity in reproductively young mice

The increase of CD38 in the ovary with age is coincident with decreased NAD+ metabolism, an altered immune landscape, and reduced reproductive potential, so we hypothesized that a reduction in CD38 may result in enhanced reproductive longevity. Therefore, we investigated reproductive function in CD38 KO mice and validated that CD38 expression was completely eliminated in the ovary (Figure 1I). Loss of ovarian follicles, and particularly primordial follicles which define the ovarian reserve or reproductive lifespan of an individual, is a primary quantitative metric of reproductive aging ^36^. Thus, we classified and quantified follicles in ovarian histological sections from both CD38 KO and WT mice across the reproductive lifespan (Figure 3A). As expected, the number of follicles per ovarian area decreased with age, and this was irrespective of follicle class (Figure 3B-F). When comparing genotypes, ovaries from CD38 KO mice exhibited a larger number of total follicles per ovarian area at reproductively young time points (2 months and 5 months) compared to WT controls (Figure 3B). This increase was largely due to an increase in primordial follicles or the ovarian reserve (Figure 3C), although there was also a tendency towards an increased number of primary, secondary, and antral follicles per ovarian area in ovaries from CD38 KO mice relative to controls. There were no differences in follicle counts between the two genotypes at 12.5 and 20 months. When examining the proportion of each follicle class across genotypes and ages, a larger fraction of the total follicle pool was composed of primordial follicles in ovaries from CD38 KO mice compared to controls at the reproductively young time points (Figure 3G).

**Figure 3:**
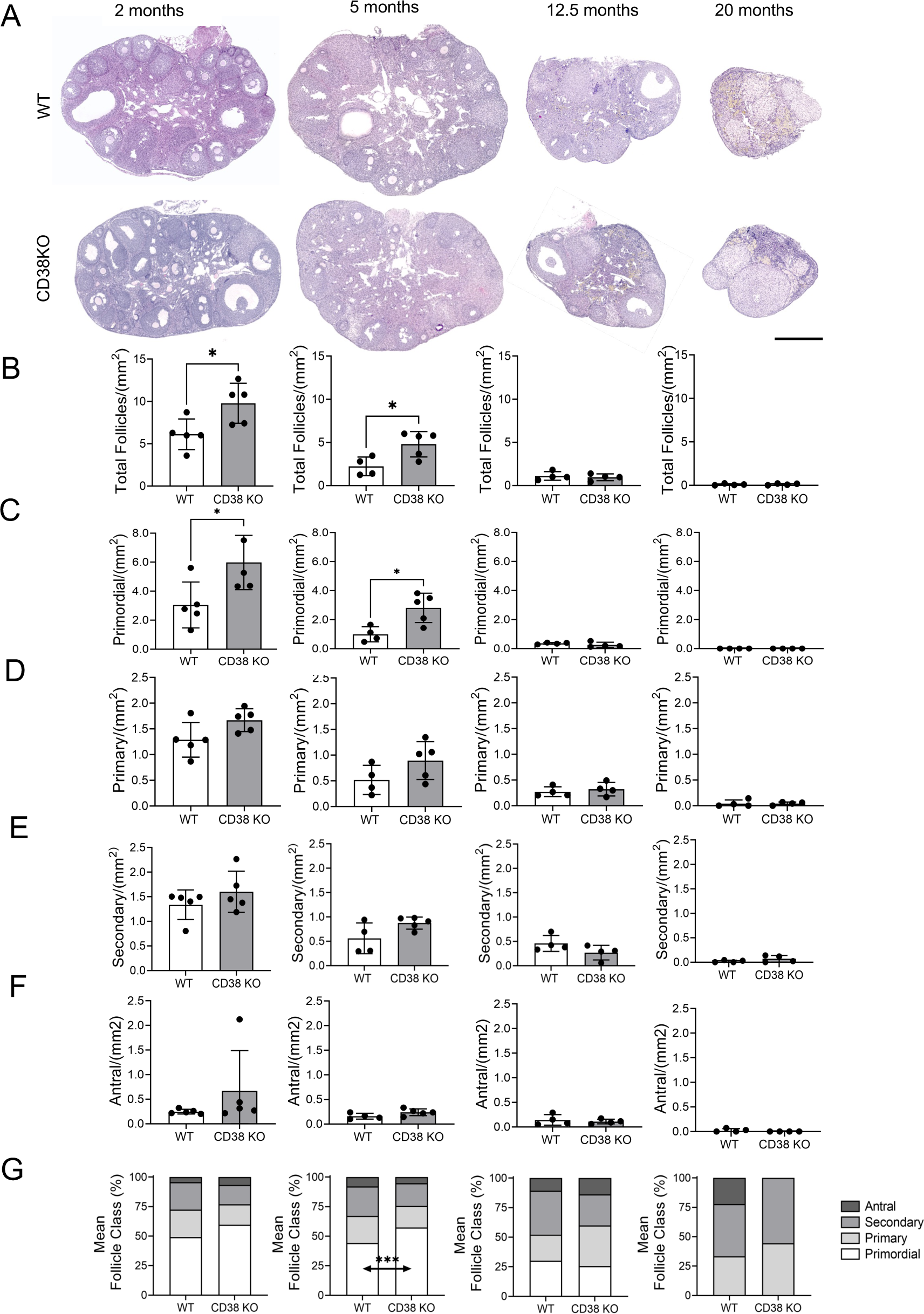
Reproductively young CD38 KO female mice have a larger ovarian reserve compared to age-matched WT. (A) Representative images of H&E stained ovarian sections from 2, 5, 12.5 and 20 months old WT and CD38 KO mice visualized by bright-field microscopy, 500 µm scale. (B) Total follicle quantification in ovaries from 2, 5, 12.5 and 20 months old WT and CD38 KO mice (n=4-5 per group). (C) Primordial, (D) primary, (E) secondary, (F) antral follicles quantification in ovaries from 2, 5, 12.5 and 20 months old C57BL/6J WT and CD38 KO mice (n=4-5 per group) (Paired t-test statistical analysis *p<0.05) (G) Follicle classes detected as a percentage of the total ovarian follicle pool in ovaries from 2, 5, 12.5 and 20 months old WT and CD38 KO mice. (Two-way ANOVA statistical analysis ***p<0.0005).

To determine whether the increase in the ovarian reserve in reproductively young CD38 KO mice translated into improved fertility, we performed a standard 6-month breeding trial with either WT and CD38 KO females at three different reproductive ages (young, mid, and advanced) and reproductively active adult WT males of proven fertility (Figure 4A). Readouts included time to achieve first litter, total litters and number of pups per dam and total number of pups per litter (Figure 4B-F). In the breeding trial, the time it took to achieve the first litter increased with age consistent with an age-dependent decline in reproductive function (Figure 4B). Interestingly, at both reproductively young and mid age timepoints, CD38 KO tended to achieve their first litter faster than age-matched WT controls, and the CD38 KO mice exhibited less variability in timing (Figure 4B). There was a prominent age dependent decline in the number of litters produced per dam irrespective of genotype (Figure 4C). At each time point, there was no difference in the number of litters produced per dam when comparing CD38 KO females to WT controls. However, at both the young and mid reproductive ages, there was a tendency towards a larger number of litters per dam in the CD38 KO mice relative to WT controls. Interestingly, although the number of litters per breeder was similar between CD38 KO animals and WT controls, the reproductively young CD38 KO females produced a significantly larger number of pups per litter during the breeding trial period (Figure 4D) and showed overall a larger number of pups per dam (Figure 4E). These data demonstrate that the increased number of follicles observed in reproductively young CD38 KO females is associated with enhanced fecundity. However, just as the increased follicle number is not sustained during aging, neither is the increase in total number of pups per dam (Figure 4E).

**Figure 4:**
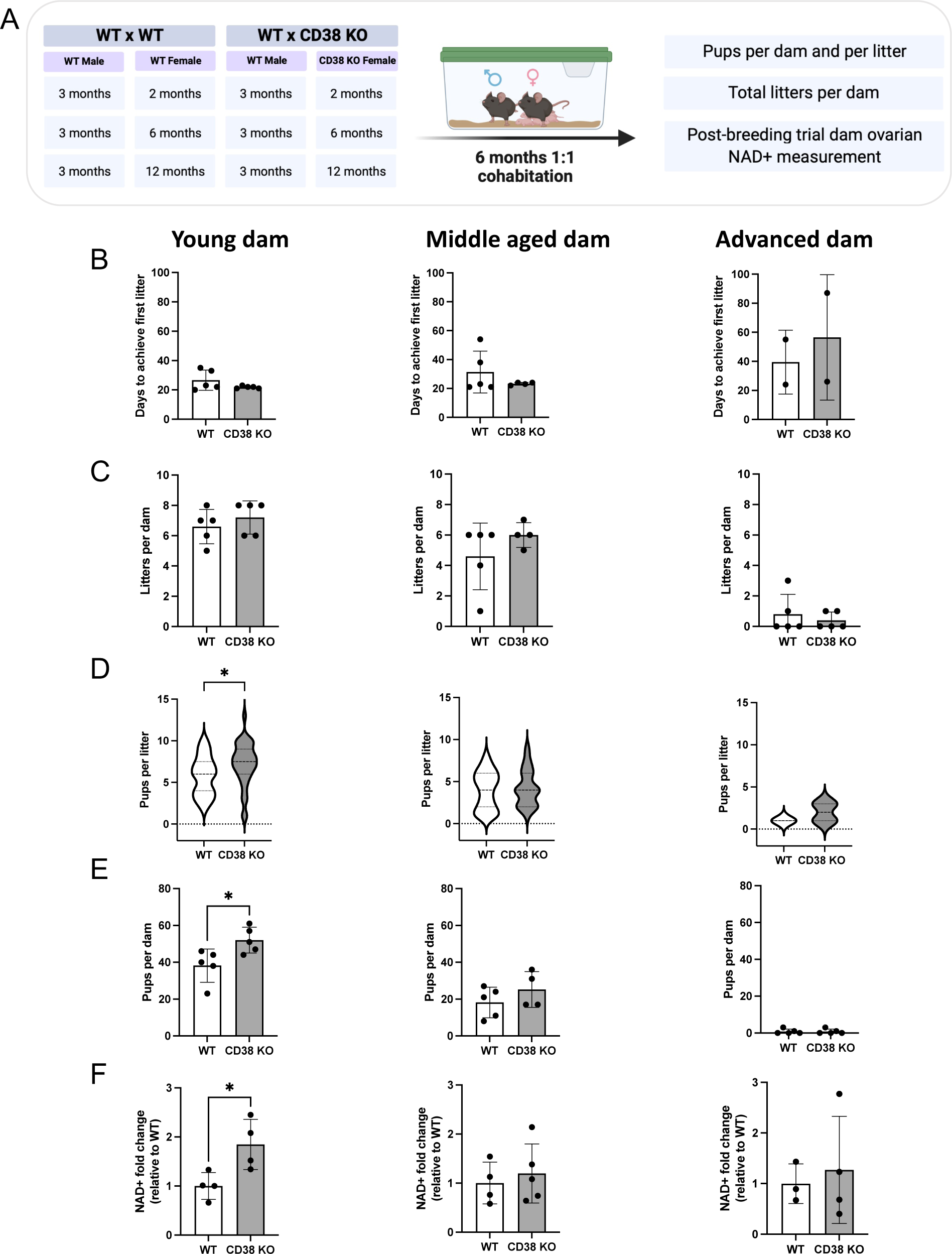
Reproductively young CD38 KO female mice generate more pups and have higher ovarian NAD+ levels compared to age-matched WT. (A) Breeding schematic of 6-months breeding trial crossing 2 (young), 6 (middle aged), 12 months (advanced) old WT and CD38 KO females with 3 months old WT males. (B) Number of days to generate their first litter and (C) total number of litters for each dam in the 3 different female age groups (young, middle-aged and advanced) throughout the 6-months breeding trial. (D) Total number of pups per litter and (E) total numbers of pups per dam generated by the 3 different female age groups (young, middle-aged and advanced) throughout the 6-months breeding trial. (Unpaired t-test statistical analysis *p<0.05) (F) Ovarian NAD+ levels of ovarian tissues collected from each female mare participating to the 6-months breeding trial (n=3-4 per group, Mann Whitney test statistical analysis *= p < 0.05)

### 6. Increased ovarian NAD+ is associated with enhanced fecundity

CD38 regulates ovarian NAD+ consumption, and NAD+ is important for reproductive function ^13^. Thus, we anticipated that the enhanced reproductive function both in terms of follicle number and fecundity in CD38 KO mice may be due to increased NAD+ levels. Thus, ovaries of the female dams were collected at the end of the breeding trial and analyzed for NAD+ content via Mass Spectrometry. NAD+ levels were significantly higher in ovaries from CD38 KO mice compared to WT controls at the reproductively young time point, although no differences between genotypes were observed at the older timepoints (Figure 4E). These data demonstrate that loss of CD38 in reproductively young females results in increased ovarian NAD+ levels which likely underlies improved reproductive outcomes.

### 7. Neonatal ovaries from CD38 knock-out mice reveals a delayed establishment of the ovarian reserve

To further investigate why reproductively young CD38 KO mice have a larger number of primordial follicles, we investigated neonatal stages of ovarian development when the ovarian reserve is established. Female mice and humans are born with a fixed number of primordial follicles, also known as the ovarian reserve, which intrinsically determines the timeline of female reproductive longevity and fertility by virtue of the size of the initial follicular reserve and the rate of follicular death or maturation ^37^. The initial formation of the ovarian reserve is a complex embryonic developmental process and it occurs during the early stages of oogenesis which involves germ cell cyst formation, organelle donation, and eventual cyst breakdown to form the primordial follicle reserve ^38^. The timeline for mouse oogenesis differs between genetic mouse strains, but the C57BL6J strain undergoes complete cyst breakdown by postnatal-day-two (P2) ^39^. CD38 protein is expressed in P2 ovaries, and its localization was completely extrafollicular similar to what we observed in adult ovaries (Figure 5A and A’). As expected, there was no CD38 expression in P2 ovaries from CD38 KO mice (Figure 5B).

**Figure 5:**
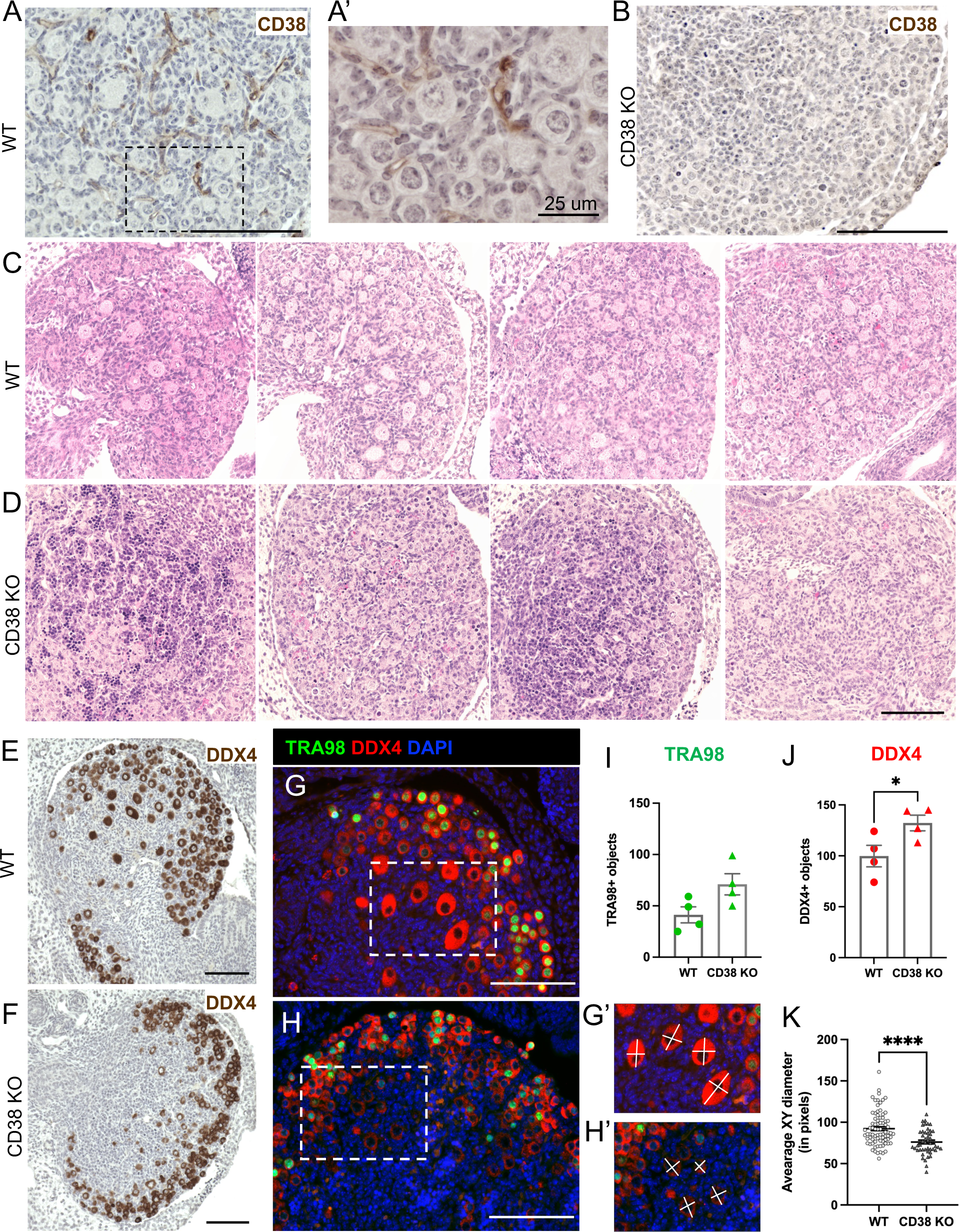
Neonatal ovaries from CD38 KO mice have delayed establishment of the ovarian follicular reserve. (A) Representative immunohistochemistry images of postnatal day 2 mouse ovaries from WT animal strained with CD38, 100 µmscale (A’) Higher magnification images of 5A, 25 µm scale (B) Representative immunohistochemistry images of postnatal day 2 mouse ovaries from CD38 KO animal strained with CD38 as control, 100 µm scale. Representative images of H&E stained ovarian sections from postnatal day 2 (C) WT (n=4) and (D) CD38 KO (n=4) mice visualized by bright-field microscopy, 100 µm scale. Representative immunohistochemistry images of postnatal day 2 mouse ovaries from (E) WT and (F) CD38 KO animals stained with the germ cell marker DDX4, 100 µm scale. Representative immunofluorescent images of postnatal day 2 mouse ovaries stained with TRA98 (in green) and DDX4 (in red) antibodies to label germ cells in (G) WT and (H) CD38 KO, 100 µm scale. Dashed rectangles in G and H show the borders of zoom in (G’) and (H’), respectively. (I) Quantification of TRA98 positive germ cells in WT and CD38 KO postnatal day 2 ovaries (n=4 for each genotype) (Unpaired t-test statistical analysis p= 0.0614). (J) Quantification of DDX4 positive germ cells in WT and CD38 KO postnatal day 2 ovaries (n=4 for each genotype) (Unpaired t-test statistical analysis *= p < 0.05). The white lines in (G’) and (H’) show examples of measurements for calculating the average XY diameter of DDX4+ germ cells (K) Average XY diameter of DDX4+ germ cells in the medullary region of WT and CD38KO postnatal day 2 ovaries (n=64 and n= 41 DDX4+ cells analyzed from total of four WT and CD38 KO ovary sections, respectively (Unpaired t-test statistical analysis ****p<0.0001).

When comparing the histology of P2 ovaries from CD38 KO and WT mice, we observed strikingly different morphology between genotypes (Figures 5C-D). Whereas ovaries from WT mice exhibited individualized primordial and primary follicles, ovaries from CD38 KO mice had fewer fully formed primordial follicles and contained many small clusters that were stained darkly with hematoxylin in the central region of the ovary (Figure 5C-D).

To further characterize the ovaries from both WT and CD38KO mice, we performed immunohistochemistry with the germ cell markers DDX4 (DEAD-Box Helicase 4, also known as VASA), a broad marker for germ cells from early embryonic development until they form antral follicles in adult ovaries ^40, 41^. We observed that ovaries from WT mice had larger DDX4 positive oocytes in the central medullary region of the ovary (Figure 5 E) while oocytes in CD38 KO mouse ovaries tended to be smaller and confined to the outer cortex (Figure 5F). We confirmed this observation via immunofluorescence staining for both DDX4 and TRA98 (Germ cell-specific antigen, also known as GCNA1) (Figure 5G-H). TRA98, like DDX4, is first expressed in germ cells during early embryonic development, but TRA98 expression is lost from oocytes in the medulla of the postnatal ovary when they arrest at the dictyate stage and form primordial follicles^42^. Quantification of TRA98+ and DDX4+ oocytes revealed that there was an increasing trend in TRA98+ germ cells and significantly more DDX4+ germ cells in ovaries from CD38 KO mice relative to age-matched controls (Figure 5 I and J). Moreover, there were differences in the size and distribution of the germ cells between the two genotypes. The average diameter of DDX4+ oocytes from WT mice were larger than those from CD38 KO mice (Figure 5G’, H’, K). These larger oocytes in ovaries from WT mice were part of more advanced stage follicles that were developing in the central medullary region of the ovary (Figure 5C, E, G). In contrast, the smaller oocytes in ovaries from CD38 KO mice tended to be confined to the outer cortex (Figure 5D, F, H). Overall, our analysis revealed that ovaries from WT mice had an increased number of more advanced stage oocytes at P2 relative to CD38 KO mice. These data suggest that CD38 KO mice may have an extended window of follicle formation during early development which may enable the endowment of a larger ovarian reserve.

## Discussion

The ovary is the first organ to age contributing to infertility and loss of endocrine function. NAD+ metabolism is involved in energy metabolism, DNA repair, and redox signaling, and NAD+ decline in whole ovarian tissue and oocytes occurs with advanced reproductive age ^8, 10, 11^. Supplementation of reproductively old mice with NAD+ precursors increases ovarian NAD+ levels and rescues fertility ^10^. Furthermore, NAD+-dependent proteins such as sirtuins have beneficial effects in preserving ovarian reserve and reversing ovarian aging in reproductively old mice ^10, 43–45^. Despite the importance of NAD+ metabolism in ovarian aging, the underlying mechanisms driving NAD+ decline have been elusive. In this study, we demonstrated that CD38 is a causal factor driving ovarian NAD+ decline, and that CD38 deficiency improves ovarian function early in the reproductive lifespan.

CD38 is a major NAD+-consuming enzyme in several aging tissues, including liver, adipose, spleen and skeletal muscle, with overall elevated CD38 expression observed at advanced chronologic age (24 to 32 months) ^20, 26^. Interestingly, we observed an age-dependent increase in CD38 in the ovary, coincident with a declining trend in ovarian NAD+ levels by 12 months, which represents a reproductive aging time point. Thus, our data suggest that increased CD38 levels are a hallmark of general tissue-specific aging irrespective of the chronological time frame.

CD38 degrades NAD+ to produce NAM and ADPR through its hydrolase activity, and cyclic ADPR (cADPR) via its cyclase activity, and our results demonstrate that CD38 actively regulates ovarian NAD+ metabolism ^46^. Ovaries from CD38 KO animals exhibited an increase in NAD+ levels and a concomitant decrease in NAM and ADPR levels, consistent with the substrate-product stoichiometry of enzymatic activity. Under normal physiological circumstances, NAM is constantly recycled to form NAD+ through the salvage pathway. Indeed, ovarian NAM levels correlated with NAD+ levels, and the absence of NAM accumulation indicated active biosynthesis through the salvage pathway. Moreover, at 20 months of age, when NAD+ levels in the CD38 KO ovaries were similar to WT levels, NAM levels also returned to WT levels. In contrast to NAM, ADPR levels were strongly suppressed across reproductive ages, and this suppression was still prominent at 20 months of age in CD38 KO ovaries. We speculate that ADPR, being a secondary messenger that mobilizes Ca^2+^ ions, was consistently used at all timepoints, and the observation that ADPR levels did not decrease further suggested that its levels were sufficient. We also predict that cADPR, another secondary messenger produced by CD38’s cyclase activity, would behave similarly to ADPR, but due to its low abundance, we could not robustly measure this metabolite.

Although CD38 has been implicated as an immune cell marker in the context of ovarian cancer, its role in ovarian physiology and aging has not been elucidated ^47–49^. CD38 expression was enriched in extrafollicular ovarian compartments such as the stroma, vasculature, and CLs, with minimal to no expression within follicles at any stage of development. Our analysis revealed an age-associated decrease in NAD+ levels in whole ovaries which include both follicles and extrafollicular structures. Previous reports have demonstrated that NAD+ levels decline within oocytes during reproductive aging in mice and that biochemical processes in the cumulus cells can affect the redox status of oocytes in vitro ^10 50^. Because CD38 is not expressed in the follicle, it is possible that other yet-to-be-identified factors may underlie the age-dependent decline of NAD+ in oocytes. Alternatively, there may be an indirect effect given the sophisticated system of communication and signaling between cells of the follicle-including oocytes, granulosa cells, and theca cells - and cells of the surrounding stromal tissue, such as fibroblasts, endothelial cells, and immune cells ^28^.

Although CD38 is expressed in several cell populations, it is predominantly expressed in ovarian immune cells, consistent with high CD38 expression in immune cells in other tissues ^23, 31, 51^. The localization of CD38 positive cells within the stroma and the CLs also confirms cellular identity given that these structures are enriched in leukocytes and are sites of immune infiltration and inflammation ^28, 52^. Within the ovary, immune cells are localized throughout the stroma and around the theca vasculature and play a key role in physiological ovarian processes such as ovulation. In the CLs, immune cell infiltration is essential for postovulatory events such as CL regression ^52^. Our immunophenotypic analysis revealed that CD38 is highly expressed in populations of both the innate and adaptive ovarian immune landscape, including B cells, T cells, macrophages, NK, NKT and ILCs. Resident macrophages and DC represent the most abundant immune cell type in the ovary irrespective of age. Ovarian macrophages mostly localize in the stroma, CLs, and atretic follicles contributing to CLs regression and tissue homeostasis ^24^. Macrophages also associate with growing follicles, promoting development and survival ^53^. The most represented subpopulation of T cells was the Double Negative (DN) population, characterized by expression of the alpha-beta T-cell receptor (TCR), but not the CD4 nor the CD8 co-receptors. The frequency of this subpopulation is usually very low in peripheral blood and tissues and emerging evidence suggests a key role in pathophysiological processes associated with autoimmunity and inflammation ^54^.

Ovarian immune cell composition varies across the reproductive cycle and with age ^25, 27^. Our analysis confirmed that reproductive aging impacts the ovarian immune cell landscape with a decrease in macrophages and DC and an increase in ILCs and T cells, mostly DN T cells. Similar trends in WT mouse ovaries were previously observed ^27, 32^. The decrease in innate immune cells, in particular macrophages, reflects the age-associated decrease in ovulation and wound healing and associated inflammation ^55^.

Although ovaries from CD38 KO mice have an overall similar immune composition relative to WT controls, they did exhibit unique age-related immune trends. For instance, while ovaries from WT mice had an overall increase in immune cells with age, the number remained constant in ovaries from CD38 KO mice. Moreover, ovaries from WT mice demonstrated an age-dependent decrease in monocytes and granulocytes and an increase in ILCs. However, no differences were observed in the absence of CD38, suggesting a protective effect. A hallmark of the aging ovary is accumulation of a unique population of multinucleated macrophage giant cells (MNGCs) which are associated with chronic inflammation. The absence of CD38 is associated with a slight decrease in ovarian MNGCs at 5 months of age but a significant increase at 28 months, suggesting an increased inflammatory milieu. Our data suggest an antagonistic pleiotropy immuno-modulatory role in the absence of CD38, with protective effects early in the reproductive lifespan and detrimental effects in post-fertile ages. This observation is particularly relevant given that ovarian aging, as a result of continuous follicle atresia and clearance as well as repeated cycles of ovulation-associated inflammation, wound repair and remodeling, is associated with increased ovarian inflammation, fibrosis, and stiffness ^35, 56–58^. This microenvironment has potential ramifications for pathologies such as ovarian cancer ^59–61^. To our knowledge, this is the first report showing the effect of CD38 genetic ablation in shaping the immune landscape during aging.

Genetic ablation of CD38 enhanced reproductive function in females. CD38 KO animals had more primordial follicles than wild type counterparts at 5 months of age. This increased ovarian reserve was associated with increased fecundity and ovarian NAD+. Previous reports have linked improved fertility to restoration of ovarian NAD+ levels, albeit in reproductively aged females ^10, 11^. The beneficial effect of NAD+ in ovaries can be explained by regulation of sirtuins, which depend on NAD+. For example, ovarian SIRT1 prevents follicular activation and loss, influencing key cellular processes such as autophagy and mTOR-related metabolic pathways ^43, 44, 62^. Interestingly, treatment with a SIRT1 agonist improves ovarian function in a mouse model of premature ovarian insufficiency (POI) ^63^. SIRT3 is a mitochondrial sirtuin and its ovarian expression decreases with age leading to mitochondrial dysfunction and deleterious spindle assembly ^64^. Overexpression of SIRT2 improves fertility and recapitulates the benefits of NMN supplementation ^10^.

Our data reveal an unexpected and novel role for CD38 in early follicle development. Ovaries from CD38 KO mice showed a prolonged period of follicle formation which likely underlies the larger ovarian reserve until young adulthood. At postnatal day 2, ovaries from CD38 KO mice had a larger number of germ cells that were in an earlier stage of development as evidenced by their smaller size relative to WT controls. Both extrinsic and intrinsic factors can contribute to the timeline of cyst breakdown and follicle formation during oogenesis, including starvation, exposure to endocrine disruptors and hormones, as well as genetic regulators ^65, 66^. Future studies are warranted to determine CD38-driven mechanisms regulating follicle formation.

The positive effects of CD38 deficiency on ovarian function were not sustained with age, and this may be due to several reasons. First, there could be a detrimental effect of CD38 deficiency in immune response with progressive aging. CD38 is necessary for immune cell infiltration and cell signaling, and the whole-body CD38 KO may have broader immunomodulatory effects not captured through our analysis which focused on composition rather than function. Lack of CD38 impacts immune response to infections, resulting in lower innate immune infiltration and neutrophil number ^51^. The increased accumulation of MNGCs in ovaries from CD38 mice at 28 months of age supports this possibility. Second, compensatory metabolic mechanisms in the total body CD38 KO model could override the beneficial effect of CD38 deficiency later in reproductive life. Since this phenotype can potentially be associated with the systemic lack of CD38 in the total body knockout mice, further studies with selective CD38 small molecule inhibitors or conditional knockout animal models will be informative.

Overall, our findings demonstrate that CD38 loss-of-function early in the reproductive lifespan increased NAD+ levels and improved ovarian function. However, this beneficial effect is not sustained during reproductive aging. Further studies are needed to fully understand the mechanisms underlying the CD38-NAD+ axis in the ovary in the context of age-related ovarian dysfunction. Furthermore, impaired ovarian NAD+ metabolism has been linked to various ovarian dysfunctions such as infertility, premature ovarian failure, and polycystic ovarian syndrome ^10, 67, 68^. Thus, understanding the role of CD38 in ovarian NAD+ metabolism has the potential to lead to novel therapeutic interventions for reproductive health and longevity.

## Methods

### Animals

Male and female C57BL/6J and CD38 KO on a C57BL/6J background mice were obtained from the Jackson Laboratory (Bar Harbor, ME). Mice were bred and maintained at the Buck Institute vivariµm with a 12-hour light and dark cycle on a standard chow diet. Mice were provided food and water ad libitum. Animal euthanasia was performed either as a humane endpoint or according to experimental endpoints. All procedures were performed in accordance with federal guidelines and those set forth by the Buck Institute Institutional Animal Care and Use Committees (IACUC). Ovaries were isolated from the reproductive tract of WT or CD38 KO mice aged at postnatal day 2, or 2-, 5-, 6-, 12-, 12.5- and 20-months old and either flash frozen in liquid nitrogen or placed in fixative for downstream analyses.

### Ovarian tissue fixation, processing, and embedding

Isolated ovaries were fixed in Modified Davidson’s Fixative (Electron Microscopy Sciences, Hatfield, PA, 64133-50) for two hours at room temperature and then at 4℃ overnight with gentle agitation. Tissues were then washed in 70% ethanol (IBI Scientific, Dubuque, IA., IB 15727) three times for 10 minutes each. Tissues were stored in 70% ethanol at 4°C for up to one week until they were processed for paraffin embedding. All paraffin processing was performed by the Buck Institute’s Morphology Core using an automatic benchtop tissue processor (Leica Biosystems, Wetzlar, Germany, TP1020). Paraffin-infused ovarian tissues were embedded in paraffin blocks, and tissues were sectioned at 5 micron thickness using a Leica microtome for downstream applications (Leica Biosystems, RM2155).

### RNA in situ hybridization and transcript quantification

The RNAScope 2.5 HD Red assay kit (Advanced Cell Diagnostics, Hayward, CA, Cat No. 322350) was used to detect Cd38 mRNA transcripts in ovarian sections from n=4 reproductively young (2-month-old) C57BL6/J mice. All probes, buffers, blocking and amplification reagents were provided by Advanced Cell Diagnostics. Control probes included Ppib (Cat No. 313911) as a positive control and DapB (Cat No. 310043) as a negative control. The experimental probe was specific to base pair 2-926 of the spliced Cd38 mRNA transcript (Cat No. 513061). Control probes were validated first on control slides containing mouse T3T cells provided by Advanced Cell Diagnostics and all probes were subsequently validated in C57BL6/J liver sections where Cd38 has been previously established. The experiment was performed as previously described ^58^ with the following modifications. In brief, FFPE tissues are 5 µm sections that were deparaffinized in Citrisolv (Fisher Scientific, 22-143-975) followed by dehydration in a graded ethanol series. Tissue sections were incubated in 1X RNAscope Target Retrieval buffer at 100°C for 15 minutes followed by a wash in deionized water. Sections were immediately treated with RNAscope Protease Plus and incubated for 23 minutes at 40°C in the HybEZ hybridization oven (ACD). Probe hybridization and amplification were performed according to manufacturer instructions ^69^. Sections were then rehydrated in an ethanol series and counterstained for 2 minutes in 50% Harris Hematoxylin (Fisher Scientific; Waltham, MA, 23 245651) with brief exposure to 0.02% ammonia water (Sigma-Aldrich, St. Louis, MO, 221228). Sections were imaged in brightfield using a 20X objective on a ZEISS Axioscan7 Imaging system (ZEISS, Oberkochen, Germany). Cd38 transcripts were quantified and normalized to structure area using Fiji ImageJ ^70^ in specific ovarian sub-compartments, including all follicle classes (primordial, primary, secondary, and antral), corpora lutea, the stroma, vasculature, and the ovarian surface epithelium. In brief, ovarian sub-compartments were identified and defined based on characteristic histologic morphology, and regions of interest (ROI) were drawn and measured using the Freehand selection tool. A color deconvolution macro was written to separate the 2.5 HD Red chromogen from the hematoxylin counterstain in all sections. A standard threshold value was established based on a clear Cd38 transcript signal and was used for all measurements. The calculated Cd38 transcript area was then normalized to the sub-compartment specific area to generate Cd38 transcript signal density in each ROI. A single ovarian section from each animal was analyzed and one of each ovary specific compartment was quantified per section.

### Colorimetric immunohistochemistry (IHC)

We performed colorimetric IHC to detect CD38 (Abcam, Cambridge, UK, ab216343 1:1000 for 2-month-old ovary and 1:200 for P2 ovary) and DDX4 (Abcam, ab270534 1:2000). Slides were deparaffinized in Citrisolv Clearing Agent (Fisher Scientific, 22-143-975) or xylene (VWR, Randor, VA, 1330-20-7) and rehydrated in a series of graded ethanol washes (100%, 95%, 85%, 70%, and 50%) that were diluted from 100% ethanol (VWR, 71002-512). Antigen retrieval was performed using an Antigen Unmasking Solution according to the manufacturer’s instructions (Vector Laboratories, Newark, Ca, H-3301-250). Slides were washed with 1X Tris-buffered saline (TBS) (Cell Signaling Technology, Danvers, MA, 12498) with 0.1% Tween-20 (Sigma-Aldrich, P1379-1L) (TBST) twice for 15 minutes and then incubated in BLOXALL Endogenous Blocking Solution, Peroxidase and Alkaline Phosphatase (Vector Laboratories, SP-6000-100) for 15 minutes at room temperature. Slides were rinsed in TBS and blocked using an avidin/biotin blocking kit (Vector Laboratories, SP-2001). Slides were then rinsed in TBS and the area around the tissue was defined using a ImmEdge Hydrophobic Barrier PAP Pen (Vector Laboratories, H-4000). Slides were incubated in protein block solution (10% normal goat secondary serum (Vector Laboratories, S-1000) in 3-5% Bovine Serum Albumin - Fraction V in-1X TBS for one hour (Rockland, Baltimore, MD, BSA-1000). Block was removed and slides were incubated in the respective primary antibody diluted in protein block solution for 1 hour or overnight in a humidified chamber. Slides were washed three times in TBST for 5 minutes each and then incubated with a drop of Goat Anti-Rabbit IgG Antibody (H+L) biotinylated Ready to Use Secondary, (Vector Laboratories, BP-9100-50, undiluted) for one hour at room temperature. Slides were then incubated with avidin/biotin complex (ABC) (Vector Laboratory, PK-6101) according to manufacturer’s instructions. Detection was then performed using a DAB Peroxidase (HRP) Substrate Kit (Vector Laboratories, SK-4105) according to manufacturer’s instructions. DAB substrate reaction was stopped by incubating in dH20 for 5 minutes. Slides were then subjected to a hematoxylin counterstain by incubating slides in the following solutions for described times: 2 minutes in Modified Mayer’s Hematoxylin (StatLab, McKinney, TX, HXMMHGAL), 2 washes with dH20, 1 minute incubation in 10% Acetic acid (Sigma-Aldrich, 27225-1L-R), 2 washes with dH20, 2 minutes in bluing reagent (Fisher Scientific, 23 245681), rinse in dH20, 1 minute in 70% ethanol, 1 minute in Eosin Y Stain (StatLab, STE0150), rinse in increasing grades of ethanol baths (90%, 95%, 100%), and finally cleared in two Xylene baths. Finally, slides were mounted and coverslipped. Slides were cured overnight and then brightfield images scans were taken with 20X or 40X objectives using either the ZEISS Axioscan 7 Imaging system or the Keyence All-in-One Fluorescence Microscope (Keyence, Itasca, IL, BZ-X800E).

### Immunofluorescence

We performed immunofluorescence in ovarian sections to determine co-labeling of CD38 with either F4/80 (Abcam, ab6640 1:200) or CD31 (ARP American Research Products, Waltham, MA, DIA-310 1:20). Histological sections were subject to the same deparaffinization, rehydration, antigen retrieval, and protein blocking steps as previously described. Block was removed and slides were incubated in the respective primary antibodies diluted in the protein block solution overnight in a humidified chamber. The primary antibodies were then removed and sections were washed with TBST three times for 5 minutes each. Sections were then incubated with goat Alexa 488 and goat Alexa 555 fluorophore-conjugated secondary antibodies (Invitrogen, Waltham, MA, A11034, Thermo FIsher Scientific i 1:1000) diluted in the protein block solution for one hour at room temperature in a humidified chamber. Sections were then washed three times for 5 minutes each with TBST. Tissues were then stained with 4′,6-diamidino-2-phenylindole (DAPI) (Invitrogen, D1306, 1:5000) for five minutes to visualize DNA. Sections were then washed three times for 5 minutes each with TBST. Slides were dehydrated in 70% ethanol for two minutes and incubated in Sudan Black B (SBB) to block endogenous autofluorescent lipofuscin accumulation in the ovary (Sigma-Aldrich, 50395, 1:100) for five minutes. Slides were washed with 70% ethanol to remove all SBB and subsequently washed three times with TBST for 5 minutes each. Slides were then mounted with ProLong Gold Antifade Reagent and allowed to cure overnight (Cell Signaling Technology, 9071). Slides were imaged using the Keyence Microscopy Imaging System with a 40X objective.

In an effort to access the initial primordial follicle reserve, postnatal day two ovaries were collected from CD38 KO and WT female pups and stained for Anti-GCNA1 (Tra98) (Abcam, ab82527, 1:500) and anti-DDX4/MVH (VASA) (Abcam, ab270534, 1:500) in accordance with approved IACUC euthanasia protocol. All fixing, sectioning, staining and mounting procedures were conducted in the same manner as previously described; however, the following changes were applied. Antigen retrieval was conducted using a citrate buffer (created in house from Citric Acid (Sigma-Aldrich, C-0759) and Sodium Citrate (Sigma-Aldrich, S-4641) diluted in dH20). All wash steps were completed using 1X Phosphate Buffered Saline (PBS), Powdered, Ultra Pure Grade (VWR, 97062-336) resuspended in dH20. The same secondary antibodies were used as described previously, however, a 1:200 dilution was used. Sections were imaged using the Keyence All-in-One Fluorescence Microscope at the 40x objective. Quantification of VASA-positive and TRA98-positive germ cells was performed using Imaris v9.9 (Bitplane). After converting files with Imaris converter for downstream analysis, they were imported into Surpass mode and germ cells were manually selected with the spot module. Positive germ cell numbers for VASA and TRA98 immunofluorescence labeling were imported to Microsoft Office Excel. A measurement of the XY diameter of oocytes in the medullary region of P2 ovaries was conducted with ZEN 3.0 (blue edition). From the measurement tab of the software, the line drawing option was selected. In order to measure each oocyte twice, two lines were drawn from four distant points. An average XY value was calculated for each oocyte after importing XY values (in pixels) into Microsoft Office Excel. Statistical analysis was performed using Prism software. t-test was used for analyzing VASA-positive and TRA98-positive germ cell numbers and XY diameters of oocytes between two genotypes.

### Immunoblot analysis

Previously frozen whole ovary tissue was lysed in RIPA buffer (Thermo Fisher Scientific, Waltham, MA, 89900) containing 1X Halt protease and phosphatase inhibitor cocktail (Thermo Fisher Scientific, 78447) and lysed using a handheld homogenizer. Samples were incubated on ice for 30 minutes and then were centrifuged at 4°C and 12,000 rpm for 10 minutes. The supernatant was then transferred to a pre-chilled new tube. Samples were diluted 1:10 in a RIPA cocktail buffer, and protein concentrations were determined using a Pierce BCA Protein Assay Kit (Thermo Fisher Scientific, 23225). Approximately 30 μg of ovarian lysate was combined with a 4x Laemmli Sample Buffer (Bio-Rad, Hercules, CA, 1610747) prepared with 2-Mercaptoethanol (Sigma-Aldrich, M6250-250ml) according to the manufacturer’s instructions. Samples were mixed well and denatured at 95°C for 5 minutes. A prestained protein marker (Thermo Fisher Scientific, 26617) and samples were loaded into individual wells of a 4-20% Mini-PROTEAN TGX Precast Protein Gel (Bio-Rad, 4561094). The gel was run in a running buffer (made in house from Sodium Chloride (Sigma-Aldrich, S9888-25G), Tris 1.0M Sterile Solution, pH 7.5 Ultra Pure Grade (VWR, 97062-936), and Tween 20 (Sigma-Aldrich, 9005-64-5) at 110V until the dye front ran off the gel. The gel was then placed in a 1X Turbo buffer (Bio-Rad, 10026938) and transferred to a 0.2 µm midi-size nitrocellulose membrane (Bio-Rad, 1620112) sandwiched between layers of Trans-Blot Turbo Midi-size Transfer Stacks (Bio-Rad, L002044B) via a semi-dry transfer using a Trans-Blot Turbo Transfer System (Bio-Rad, 1704150). The membrane was washed three times with a 1X Tris Buffered Saline (TBS) (Cell Signaling Technology, 12798) and Tween 20 (Sigma-Aldrich, 9005-64-5) (TBST) for 5 minutes each and then blocked for one hour in 0.5% Blotting-Grade Blocker (Bio-Rad, 1706404) diluted in TBST at room temperature. The membrane was then washed three times in the wash buffer for five minutes each and incubated overnight with the same CD38 primary antibody as previously mentioned (1:1000) diluted in 5% BSA (Rockland, BSA-1000) in TBST at 4°C. The membrane was then washed three times in the wash buffer for five minutes each and incubated for one hour at room temperature with agitation in an anti-rabbit HRP-conjugated secondary antibody (Cell Signaling Technology, 7074V, 1:1000) diluted in the wash buffer. The membrane was washed three times for five minutes each in the wash buffer. The HRP substrate (Fisher Scientific, AC2101) was then prepared and added to the membrane according to the manufacturer’s instructions, and the membrane was exposed and imaged using the Azure 600 imaging system (Azure Biosystems, Dublin, CA, c600). The membrane was washed and stripped using a Thermo Scientific PLUS Western Blot Stripping Buffer Restore Plus according to the manufacturer’s instructions (Fisher Scientific, PI46430). The membrane was then blocked, washed, and re-probed with a beta actin-HRP conjugated antibody (Fisher Scientific, MA515739HRP 1:1000) as described above, and re-imaged using the same Azure imaging system. Images were then analyzed in Fiji ImageJ to quantify relative amounts of the protein bands. The values of the CD38 bands were normalized to the corresponding housekeeping β-actin bands.

### In-silico analysis of publicly available ovarian bulk RNAseq data

A previously published mouse ovaries bulk RNAseq dataset was used for in-silico analysis of changes in CD38 expression in the mouse ovary across the reproductive lifespan. The previously published dataset (Zhang et al., Reproduction 2020) was obtained and downloaded from Buck Institute GCRLE database. The list of 22,086 gene markers was assessed for genes of interest and the average number of Cd38 transcripts per ovary was determined for mice at 3, 6, 9, 12, 15, and 18 months of age (n=3).

### LCMS analysis of the ovary

Extraction of NAD+ and related metabolites from flash frozen ovaries collected from mice of different ages and at the end of the breeding trial was performed as previously reported ^71^. Briefly, each ovary was homogenized in a bullet blender, vortexed for 10 sec and incubated for 30 mins on ice. Upon centrifugation at 18,000 rcf for 10 mins, the supernatant was transferred to a new eppendorf tube and dried in a speed vacuum. The final dried pellet was resuspended in starting buffer conditions for subsequent LCMS analysis. 13C NAD+ and 13C NAM (Cambridge isotope laboratories Inc. MA, USA, CLM-10671-PK and CLM-9925-PK) were used as internal standards. Ovary extracts were analyzed with QExactive Mass Spectrometer coupled to a Vanquish LC system (Thermo Scientific). Separation of metabolites was achieved in 12 minutes using a Hypercarb column (5 mm, 100X2.1 mm column, Thermo Scientific) and a gradient run of solvent A (7.5 mM ammonium acetate with 0.05% ammonium hydroxide) and solvent B (0.05% ammonium hydroxide in acetonitrile). The gradient program was as follows: 0–1 minute, 5% B; 6 minutes, 60% B; 6.1-7.5 minutes, 90% B; 7.6-12 minutes, 5% B. Data acquisition was performed using FullMS1-ddMS2 (Top4) within a scan range of 50–750 m/z, with a resolution of 70000, AGC target of 1e6, and a maximum IT of 100 ms. Relative levels of respective metabolites in the ovary were measured using their MS1 extracted ion chromatogram and corresponding normalized AUC. Metabolite identification was confirmed with corresponding retention time and MS2 fragmentation spectra of metabolite standards run in parallel. To compute quantitative levels, an 8-point calibration curve of pooled NAD+, NAM and ADPR standards (Sigma Aldrich, N0632, 72340, A0752 respectively) was prepared, extracted, and analyzed alongside ovary extracts. Frozen ovary weights were used to normalize to represent quantitative data as pmol/mg ovary. Aged ovaries that were too small, with corresponding inaccurate weights, were excluded.

### Flow Cytometry

For flow cytometric analysis, both ovaries from each mouse were pooled and processed according to an established protocol with slight modifications ^72^. Briefly, isolated ovaries were cut into small pieces and sequentially incubated in collagenase IV (Sigma-Aldrich, NC0217889) and DNAse I (Sigma-Aldrich, 50-178-0833) for 30 minutes and 15 minutes, respectively, in a 37°C dry bath with gently mixing. Each homogenate was then passed through an Olympus Advanced Cell Strainer, 70 μm filter (Genesee Scientific, 25-376) and the resulting filtrate was subjected to three rounds of gentle washes with PBS and centrifugation, with an incubation with Ammonium-Chloride-Potassium ACK lysis buffer (Thermo Fisher Scientific, A1049201) for 5 mins between the last two washes. Ovarian cell suspensions were then seeded in 96-well V-bottom plates and incubated with LIVE/DEAD™ Fixable Blue Dead Cell Stain Kit (Invitrogen, L34962, 1:2000) and Fc Block (BioLegend, San Diego, CA, clone S17011E, 156604) for 25 minutes at 4°C. Next, samples were stained in the staining buffer (PBS containing 1% Dialyzed fetal bovine serum (Cytiva Life Sciences, Marlborough, MA, SH30079.01), 0.09% sodium azide (Sigma-Aldrich, S2002) using the following antibody panel: PD-1-BUV615 (BD OptiBuild, clone RMP1-30, 752354, 1:50); CD27-BUV737 (BD Horizon, clone LG.3A10, 612831, 1:200), Ly-6C/g-BUV805 (BD OptiBuild, clone RB6-8C5, 741920, 1:200); Class II-BUV496 (BD OptiBuild, clone 2G9, 750171, 1:200); CD21/35-BV421 (BioLegend, clone 7E9, 123422, 1:200); NK1.1-BV605 (BioLegend, clone PK136, 108753, 1:200); CD4-BV650 (BioLegend, clone GK1.5, 100469, 1:200); CD62L-BV711 (BioLegend, clone MEL-14, 104445, 1:400); CD138-BV785 (BioLegend, clone 281-2, 142534, 1:800), CD38-Pac Blue (BioLegend, clone 90, 102719, 1:400); F4/80-BV480 (BD Horizon, clone T45-2342, 565635, 1:200); CD45-BV510 (BioLegend, clone 30-F11, 103137, 1:200), CD8-FITC (BioLegend, clone 53-6.7, 100706, 1:200); CD11b-PerCP-Cy5.5 (BioLegend, clone M1/70, 101227, 1:400), CD163-PerCP-eFluor_710 (Invitrogen, clone TNKUPJ, 46-1631-80, 1:200), CD127-PE (BioLegend, clone S18006K, 158204, 1:400), CD11c-PE-Fire_810 (BioLegend, clone QA18A72, 161105, 1:200), CD19-PE-Dazzle_594 (BioLegend, clone 6D5, 115553, 1:200), CD31-PE-Cy7 (Invitrogen, clone 390, 25-0311-81, 1:200), TCRgd-APC (BioLegend, clone GL3, 118115, 1:400); IgD-Alexa_Fluor_647 (BioLegend, clone 11-26c.2a, 405707, 1:800), CD68-AF700 (BioLegend, clone FA-11, 137025, 1:200); IgM-APC-Cy7 (BioLegend, clone RMM-1, 406515, 1:200); CD3-APC-Fire_810 (BioLegend, clone 17A2, 100268, 1:200); LY6G-BUV563 (BD Horizon, clone 1A8, 612921, 1:200). Staining was done at 4°C for 30 minutes. Flow cytometry acquisition was done using a 5-laser Cytek Aurora spectral flow cytometer (Cytek Biosciences, Fremont, CA, 16UV-16V-14B-10YG-8R). Spectral Unmixing was done using Spectoflo v3.0.3 software of Cytek and further flow analysis and UMAP plots were done using Flowjo software (BD Biosciences, Franklin Lakes, NJ, V10.8.1).

### MNGC quantification

For multinucleated giant macrophage cell (MNGC) quantification, three H&E stained ovarian sections from 2, 5, 12.5, 20, and 28 month old C57BL6/J WT (n=5, 4, 4, 4, 5; respectively) and CD38 KO (n=5, 5, 5, 4, 5; respectively) mice were analyzed to compare observable temporospatial macrophage dynamics in these groups. Section images from those used for follicle classification were analyzed. The presence of MNGCs was visually determined according to previously published criteria ^33^. Quantification of the MNGC area was performed in Fiji ImageJ by separating yellow MNGC signal from the remainder of the H&E stain using a color deconvolution macro and measurement of MNGC structures using threshold adjustment. Ovarian section area was also measured using threshold adjustment and MNGC area was reported as a % of section area. Three sections were analyzed from ¼, ½, and ¾ through each serially sectioned ovary and the average of the 3 measurements was determined to generate ovarian MNGC area density for each animal.

### Follicle classification and counting

Follicle classification and counting was performed to quantify ovarian reserve and activated follicles in WT and CD38 KO ovaries across the reproductive lifespan. Serial sections from 2, 5, 12.5, 20, and 28 month old C57BL6/J WT (n=5, 4, 4, 4, 5; respectively) and CD38 KO (n=5, 5, 5, 4, 5; respectively) were stained with Hematoxylin and Eosin (H&E) according to standard protocol ^36^. Sections were imaged in brightfield using a 20X objective on a ZEISS Axioscan7 Imaging system and the EVOS FL Auto Cell 2 Imaging system (Invitrogen, Waltham, MA, AMAFD2000). Follicles were classified according to previously published criteria ^36^ and counted by class specific annotation on ZEISS ZEN 3.4 (blue edition) or EVOS FL Auto 2 Software (Rev. 2.0.2094.0). In brief, primordial follicles were classified as oocytes surrounded by an incomplete or complete layer of squamous granulosa cells. Primary follicles were characterized as oocytes surrounded by a single layer of cuboidal granulosa cells. Primordial and primary follicles were counted irrespective of whether the oocyte nucleus was visible in the section. Secondary follicles were classified as oocytes surrounded by more than one layer of cuboidal granulosa cells. Antral follicles were classified by the presence of a visible antrum and a cumulus oocyte complex surrounded by multiple layers of mural cells. Only secondary or antral follicles with an oocyte nucleus present in the section were included to avoid duplicate counting of these large later stage follicles. Only morphologically normal follicles were counted. Atretic follicles, as distinguished by granulosa cells with pyknotic nuclei and shrunken oocytes with condensed and fragmented chromatin, were not included. The area of each analyzed ovarian section was measured by using the freehand spline tool in ZEN 3.4 (blue edition) or the freehand polygon tool on the EVOS FL AUTO. Follicle counts were normalized to the sum of the areas of all analyzed ovarian sections as a measure of ovarian follicle density. The fraction of each follicle class relative to the total follicle count for each ovary was also reported.

### Breeding trial

To assess fertility, a breeding trial was performed in which CD38 KO and WT female virgin mice at 2, 6, and 12 (n= 5, 4, 5) months of age were individually housed with 3-month-old WT males (n= 5, 5, 5) of proven fertility for a continuous 6-month period. One breeding pair from the mid CD38 KO female x WT male group generated no pups throughout the duration of the breeding trial and during the post-breeding trial tissue collection, the female had a large pituitary tumor. Due to this disease state, this breeding pair was excluded from the data analysis. Breeding cages were checked daily, and the number of litters, number of pups born, and pup survival were recorded. At the end of the breeding trial, we harvested and flash froze ovaries to assess NAD+.

### Hematoxylin and eosin staining

Histological sections were subjected to the same deparaffinization and rehydration as previously described. Slides were incubated with the following protocol: 2 minutes in Modified Mayer’s Hematoxylin (StatLab, McKinney, TX, HXMMHGAL), 2 washes with dH20, 1 minute incubation in 10% Acetic acid (Sigma-Aldrich, 27225-1L-R), 2 washes with dH20, 2 minutes in bluing reagent (Fisher Scientific, 23 245681), rinse in dH20, 1 minute in 70% ethanol, 1 minute in Eosin Y Stain (StatLab, STE0150), rinse in increasing grades of ethanol baths (90%, 95%, 100%), and finally cleared in two Xylene baths. Finally, slides were mounted, coverslipped, and cured overnight.

### Statistical analysis

Unpaired t-test, Paired t-test, Two-way ANOVA test and Mann–Whitney test were performed where appropriate as reported in the figure legends. All statistical analyses were performed using GraphPad Prism software (Version Version 9.5.1). A comparison was considered significant if p-values were less than 0.05.

## Acknowledgments

We thank Dr. Elena Silva, Kaitlyn Vitangcol, Stella Breslin, and Harris Ingle for technical assistance and useful discussion.

## Author contributions

Conceptualization, R.P., P.V.A.K., L.H., F.E.D. E.V.; Methodology, P.V.A.K., L.H., C.H, J.B., B.S.K., K.H.; Formal analysis, R.P., P.V.A.K., L.H., C.H, B.S.K., K.H., H.M.; Investigation, P.V.A.K., L.H., C.H, R.R., G.Z., B.S.K., K.H., L.P., R.T., G.V.H.; Writing—original draft preparation, R.P., P.V.A.K., L.H., C.H., Writing—review and editing, R.P., P.V.A.K., F.E.D.; Supervision, R.P., F.E.D., E.V.; Funding acquisition, E.V. All authors have read and agreed to the published version of the manuscript.

