## Supplemental Information for "CD38 regulates ovarian function and fecundity via NAD^+^ metabolism"

### Figure S1

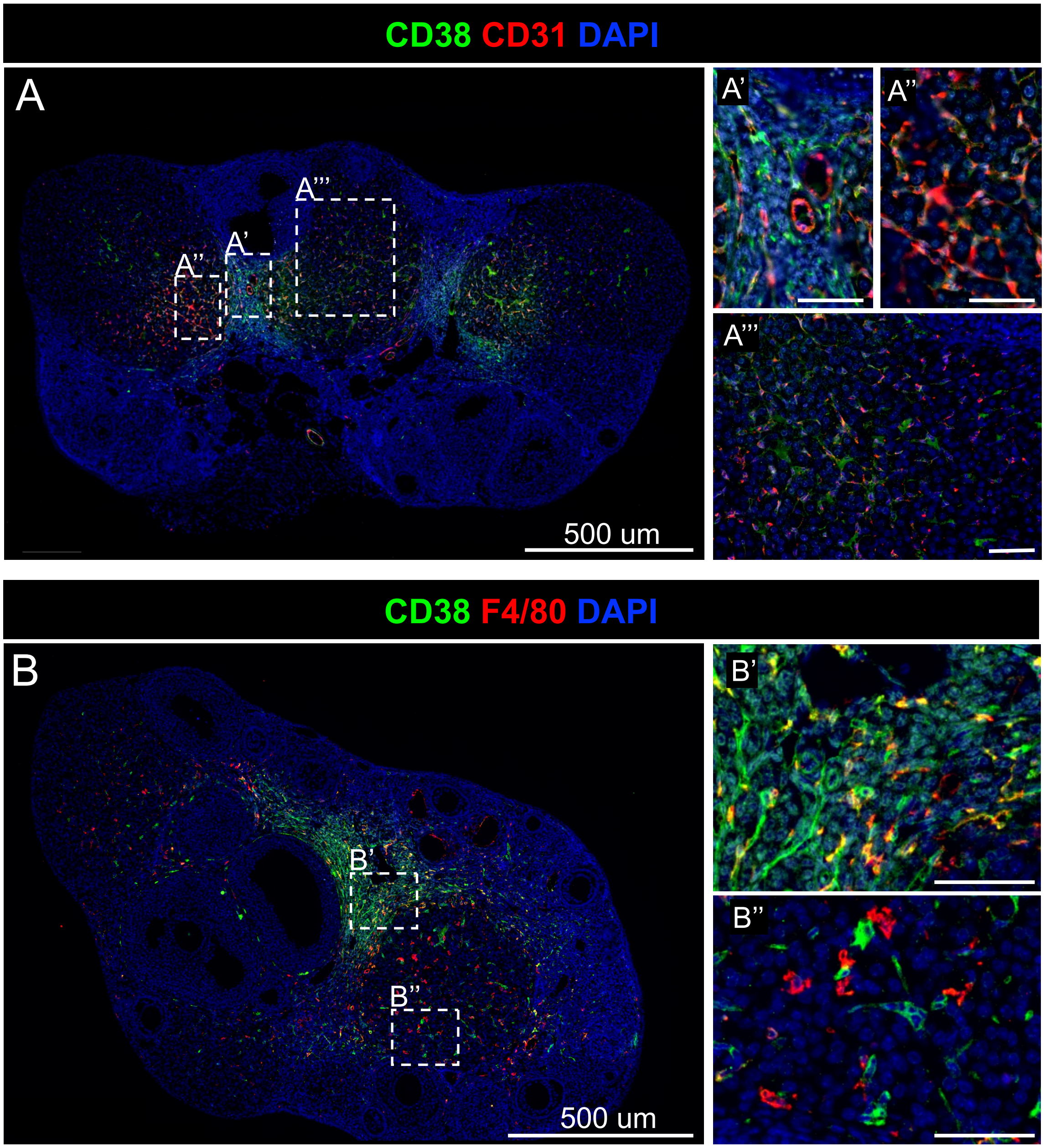

**Figure S1: Co-labeling of ovarian CD38 with macrophages and endothelial cells** (A) Immunofluorescence image of a 2-month-old WT mouse ovary showing the expression of CD38 and the endothelial cell marker CD31, 500 μm scale, (A' - A''') Higher magnification images of S2A, 50 μm scale (B) Immunofluorescence image of a 2-month-old WT mouse ovary showing the expression of CD38 and the macrophage marker F4/80, 500 μm scale (B' - B'') Higher magnification images of S2B, 50 μm scale.

### Figure S2

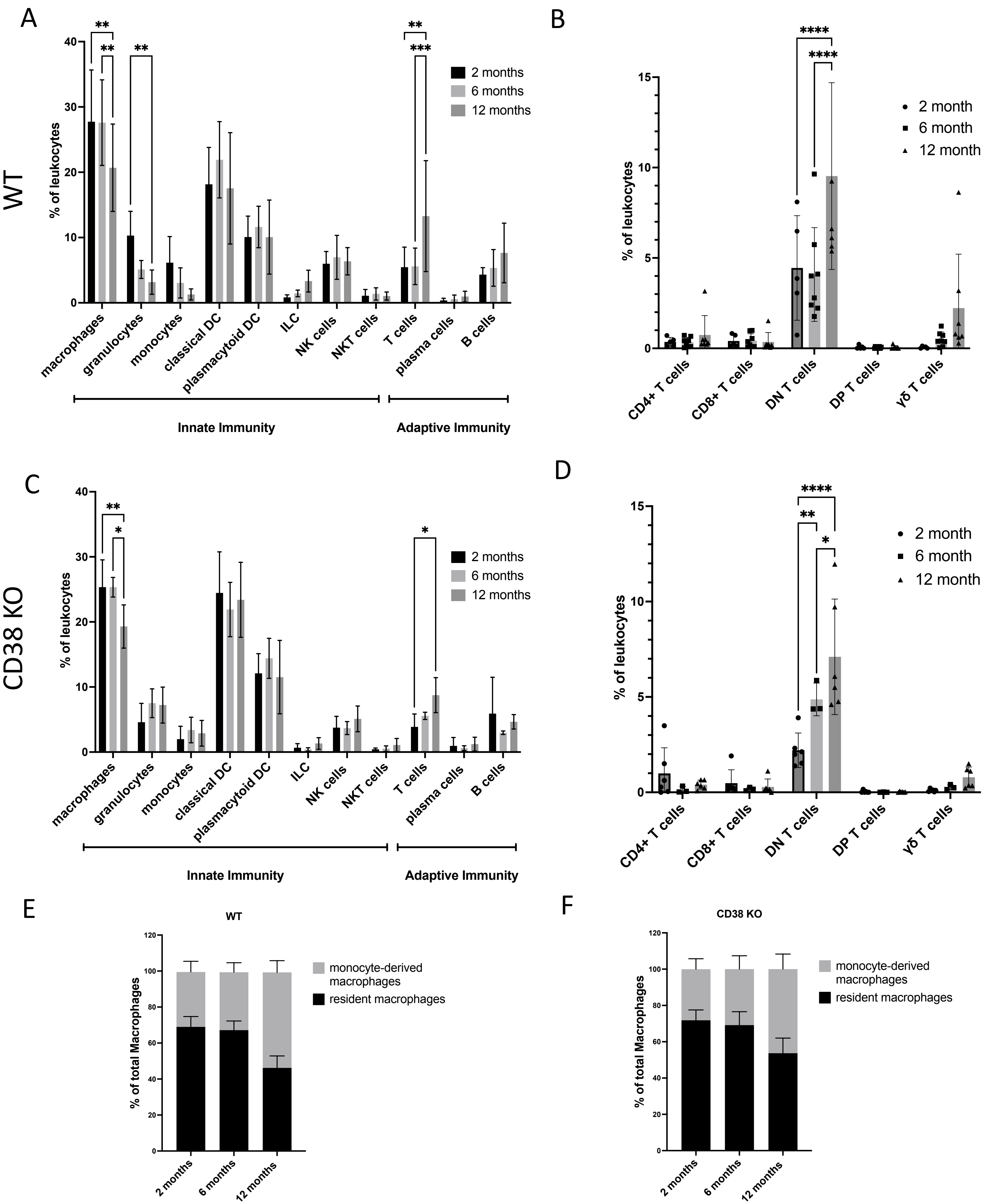

A
B
C
D
E
F
WT
CD38 KO
% of leukocytes
2 months
6 months
12 months
2 month
6 month
12 month
macrophages
granulocytes
monocytes
classical DC
plasmacytoid DC
ILC
NK cells
NKT cells
T cells
plasma cells
B cells
Innate Immunity
Adaptive Immunity
CD4+ T cells
CD8+ T cells
DN T cells
DP T cells
γδ T cells
% of total Macrophages
monocyte-derived macrophages
resident macrophages

**Figure S2: Immunophenotyping profile of WT and CD38 KO mouse ovaries during reproductive aging shows a shift from innate to adaptive immunity and a decrease in resident macrophages during reproductive aging.** (A) Percent of innate and adaptive leukocyte populations in WT ovaries throughout reproductive aging (2-12 months). (B) Percent of T cell subpopulations in WT ovaries throughout reproductive aging (2-12 months). (C) Percent of innate and adaptive leukocyte populations in CD38 KO ovaries throughout reproductive aging (2-12 months). (B) Percent of T cell subpopulations in CD38 KO ovaries throughout reproductive aging (2-12 months). (n=4-8 per group, mean ± SEM, Two-way ANOVA statistical analysis *** = $p < 0.0005$). Relative percent of monocyte-derived (CD11c+) and resident (CD11c-) macrophages within (E) WT and (F) CD38 KO ovaries throughout reproductive aging (2-12 months).

### Figure S3

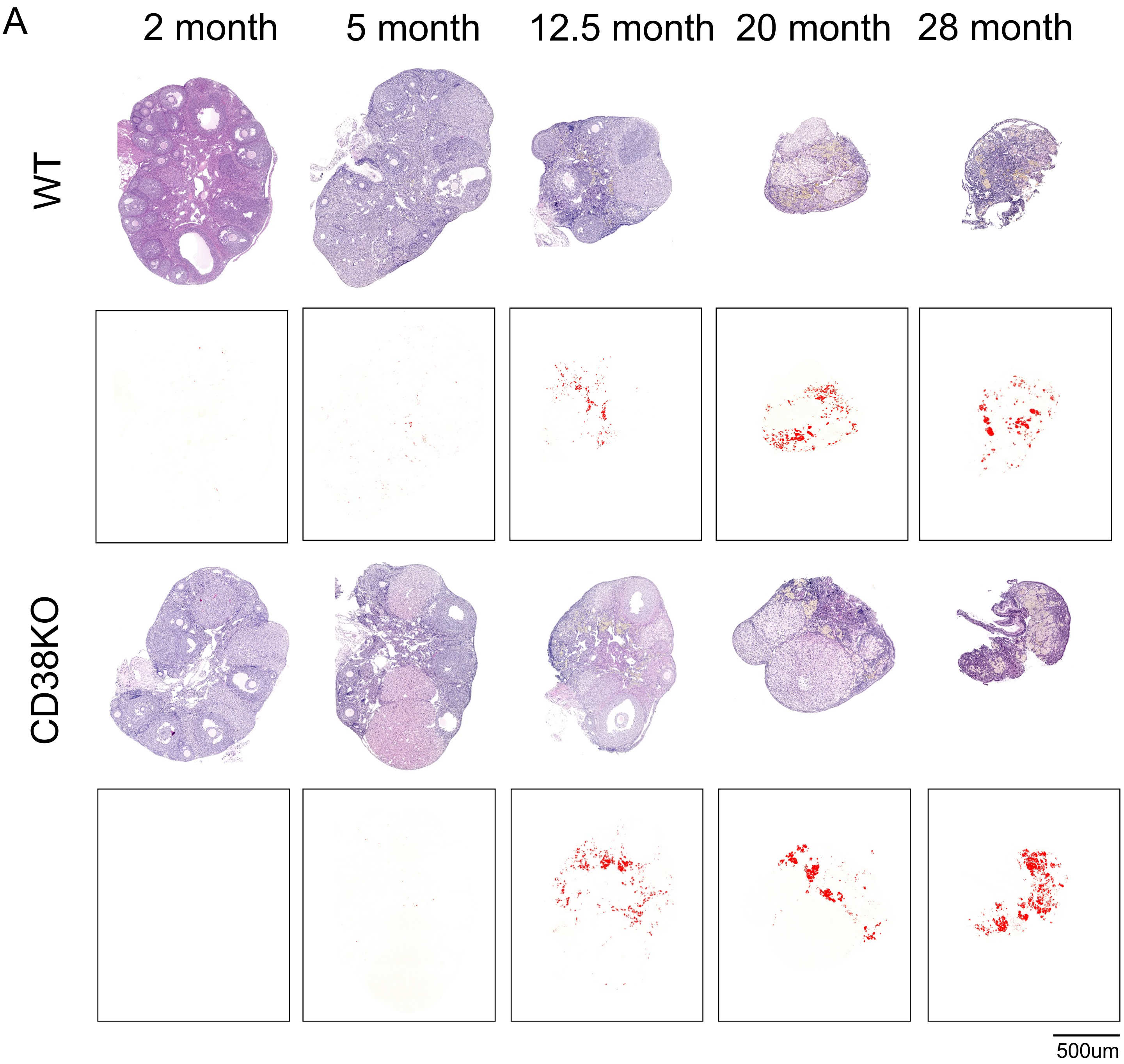


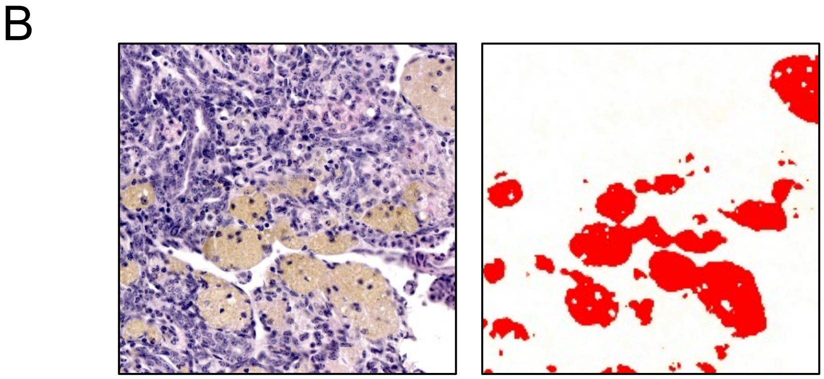

**Figure S3: Image processing and signal threshold analysis of H&E stained sections yields quantitative understanding of macrophage dynamics with age in WT and CD38 KO ovarian micro environment.** (A) Representative images of H&E stained ovarian sections from 2, 5, 12.5, 20, and 28 month old WT and CD38KO mice visualized by bright-field microscopy with corresponding images from MNGC quantification. (B) Representative image of MNGCs observed in 28 month old WT ovaries with corresponding image of thresholding used for quantification.
